# CytoTalk: *De novo* construction of signal transduction networks using single-cell RNA-Seq data

**DOI:** 10.1101/2020.03.29.014464

**Authors:** Yuxuan Hu, Tao Peng, Lin Gao, Kai Tan

## Abstract

Single-cell technology has opened the door for studying signal transduction in a complex tissue at unprecedented resolution. However, there is a lack of analytical methods for *de novo* construction of signal transduction pathways using single-cell omics data. Here we present CytoTalk, a computational method for *de novo* constructing cell type-specific signal transduction networks using single-cell RNA-Seq data. CytoTalk first constructs intracellular and intercellular gene-gene interaction networks using an information-theoretic measure between two cell types. Candidate signal transduction pathways in the integrated network are identified using the prize-collecting Steiner forest algorithm. We applied CytoTalk to a single-cell RNA-Seq data set on mouse visual cortex and evaluated predictions using high-throughput spatial transcriptomics data generated from the same tissue. Compared to published methods, genes in our inferred signaling pathways have significantly higher spatial expression correlation only in cells that are spatially closer to each other, suggesting improved accuracy of CytoTalk. Furthermore, using single-cell RNA-Seq data with receptor gene perturbation, we found that predicted pathways are enriched for differentially expressed genes between the receptor knockout and wild type cells, further validating the accuracy of CytoTalk. In summary, CytoTalk enables *de novo* construction of signal transduction pathways and facilitates comparative analysis of these pathways across tissues and conditions.

## Introduction

Single-cell RNA sequencing (scRNA-Seq) technologies are increasingly being used to characterize the heterogeneity of a complex tissue. Beyond cataloguing cell types and transcript abundance, it is critical to understand how different cell types interact with one another to give rise to the emergent tissue complexity. Signal transduction is the primary mechanism for cell-cell communication. scRNA-Seq technology holds great promise for studying cell-cell communication at much higher resolution. Using scRNA-Seq data, several methods have been developed to infer ligand-receptor pairs that are active between two cell types. Skelly et al. ^1^ and Kumar et al. ^2^ predict ligand-receptor pairs if both genes are expressed in the cell types considered. Zhou et al. ^3^ and Vento-Tormo et al. ^4^ identify ligand-receptor pairs whose expression is specific to the cell types considered. Signaling pathways are highly dynamic and crosstalk among them is prevalent. Due to these two features, simply examining expression levels of ligand and receptor genes cannot reliably capture the overall activities of signaling pathways and interactions among them ^5,6^. As a step forward, Wang et al. ^7^ developed SoptSC and Browaeys et al. ^8^ developed NicheNet to identify both ligand-receptor pairs and genes downstream of them. However, these methods are based on known annotations of signaling pathways. To our knowledge, currently no method exists to perform *de novo* prediction of the entire signal transduction pathways emanating from the ligand-receptor pairs.

Here we describe the CytoTalk algorithm for *de novo* construction of signaling network (union of multiple signaling pathways) between two cell types using scRNA-Seq data. The algorithm first constructs an integrated network consisting of intra-cellular and inter-cellular functional gene interactions. It then identifies the signaling network by solving a prize-collecting Steiner forest problem. We demonstrate the performance of the algorithm using high throughput spatial transcriptomics data and scRNA-Seq data with perturbation to the receptor genes in a signaling pathway. A software package implementing the CytoTalk algorithm has been deposited at GitHub (https://github.com/tanlabcode/CytoTalk).

## Results

### Wiring of signaling pathways is highly cell type-dependent

A hallmark of signal transduction pathways is their high level of cell-type specific wiring pattern. Single-cell transcriptome data allows us to examine the cell type-specific activity of individual signaling pathways beyond just ligand and receptor genes. To this end, we examined the canonical fibroblast growth factor receptor 2 (FGFR2) signaling pathway in two tissue types, mammary gland and skin. Four canonical downstream pathways are known to signal from FGFR2 ^9^, including Janus kinase and signal transducer and activator transcription proteins (JAK-STAT), protein kinase C (PKC), mitogen-activated protein kinase (MAPK) and phosphoinositide 3-kinase and protein kinase B (PI3K/AKT) pathways. For mammary gland, we studied FGFR2 signaling between fibroblasts and luminal epithelial cells. For skin, we studied FGFR2 signaling between keratinocyte stem cells and basal cells. Using published scRNA-Seq data ^10^ for each tissue type, we computed an expression specificity score, preferential expression measure (*PEM*) ^20,21^, for each pathway gene in each involved cell type (Fig. 1a and 1b). We found that the four canonical sub-pathways downstream of the same receptor (FGFR2) exhibit striking cell type-specific activities, depending on the cell types involved. The PI3K/AKT pathway is most active for signaling between fibroblasts and luminal epithelial cells in the mammary gland. In contrast, The JAK-STAT pathway is most active for signaling between keratinocyte stem cells and basal cells in skin. To evaluate the extent of cell type-specific wiring of signaling pathways, we examined all manually annotated signaling pathways in the Reactome database ^11^. For each pathway, we computed its cell type-specific activity score using the same mammary gland and skin scRNA-Seq data sets ^10^. We found that the majority of pathways exhibit high degree of cell type-specific activities (Fig. 1c and 1d). This is true even for the same cell types but located in different tissues, such as basal cells in mammary gland versus in skin (Fig. 1e). In summary, these results highlight the need for analytical tools for *de novo* construction of complete signaling pathways (instead of ligand-receptor pairs) using single-cell transcriptome data.

**Fig. 1.**
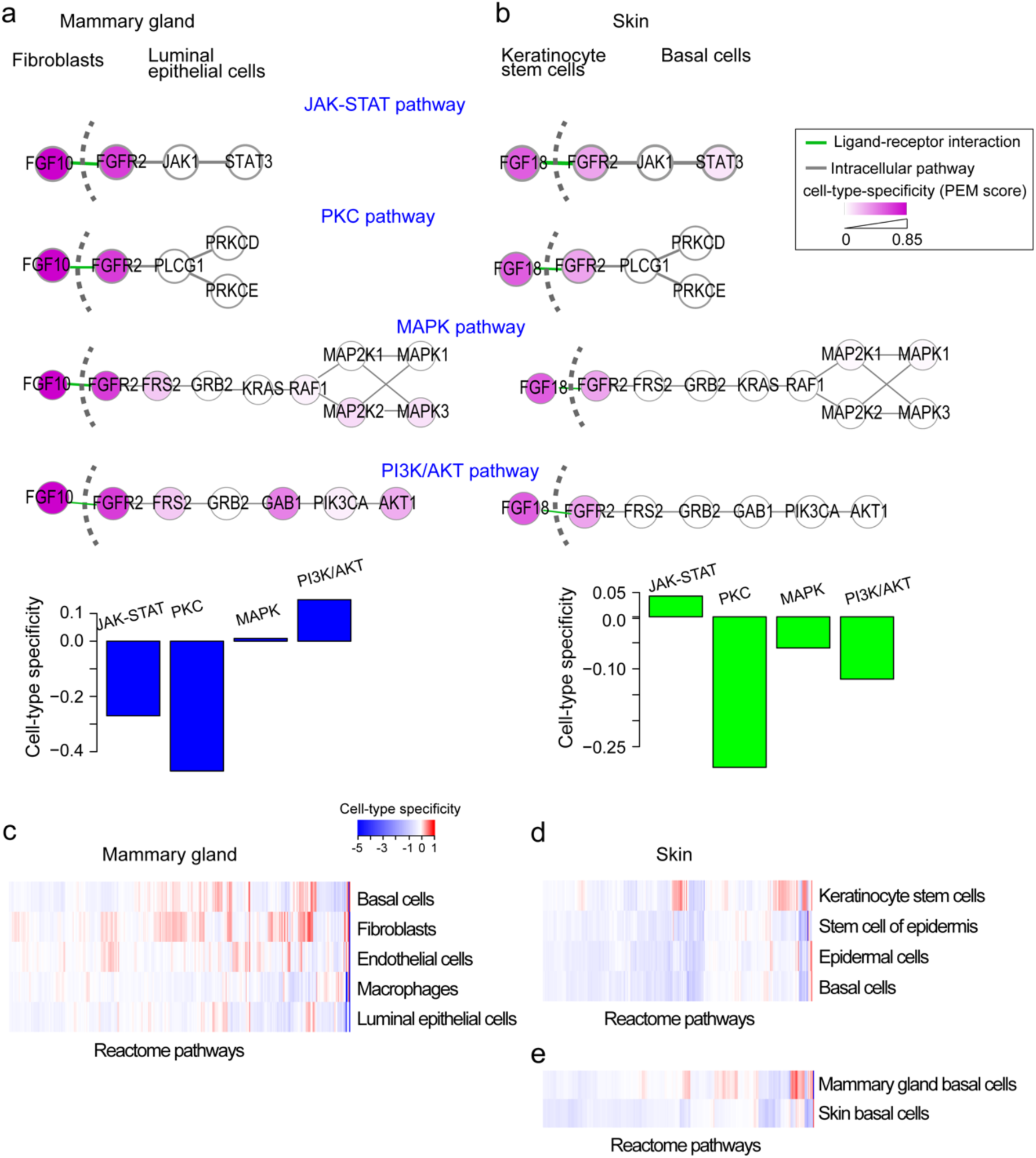
Wiring of signaling pathways is highly cell-type dependent. **(a, b)** Cell-type-specific activity of sub-pathways downstream of the fibroblast growth factor receptor 2 (FGFR2) between fibroblasts and luminal epithelial cells in mouse mammary gland **(a)** and between keratinocyte stem cells and basal cells in mouse skin **(b)**. Cell-type-specific activities of four canonical sub-pathways downstream of FGFR2 are shown. PEM, cell-type-specific activity score. Color shade of each gene node is proportional to PEM score. Top, individual pathway activities. Down, quantification of average PEM score of sub-pathway genes. (**c**) Cell-type-specific activity of Reactome pathways across five cell types in mammary gland. Pathway cell-type-specific activities were calculated using single-cell RNA-Seq data. (**d**) Cell-type-specific activity of Reactome pathways across four cell types in skin. **(e)** Differential pathway activity in basal cells from two different tissues, mammary gland and skin.

### Overview of the CytoTalk algorithm

CytoTalk is designed for *de novo* construction of a signal transduction network between two cell types (Fig. 2, Methods), which is defined as the union of multiple signal transduction pathways. It first constructs a weighted integrated gene network comprised of both intracellular and intercellular functional gene-gene interactions. Intracellular functional gene interactions are computed and weighted using mutual information between two genes. Two intracellular networks are connected via crosstalk edges (i.e. known ligand-receptor interactions). Ligand-receptor pairs with higher cell-type-specific gene expression but lower correlated expression within the same cell type (thus more likely to be involved in crosstalk instead of self talk) are assigned higher crosstalk weights. Nodes in the integrated network are weighted by a combination of their cell-type-specific gene expression and closeness to the ligand/receptor genes in the network. We use a network propagation procedure to determine the closeness of a gene to the ligand/receptor gene. With the integrated network as the input, we formulate the identification of signaling network as a prize-collecting Steiner forest (PCSF) problem ^12,13^. The rationale for using the PCSF algorithm is to find an optimal subnetwork that includes genes with high level of cell-type-specific expression and close connections to high-scoring ligand-receptor pairs. This optimal subnetwork is defined as the signaling network between the two cell types. The statistical significance of the candidate signaling network is computed using a null score distribution of signaling networks generated using degree-preserving randomized networks.

**Fig. 2.**
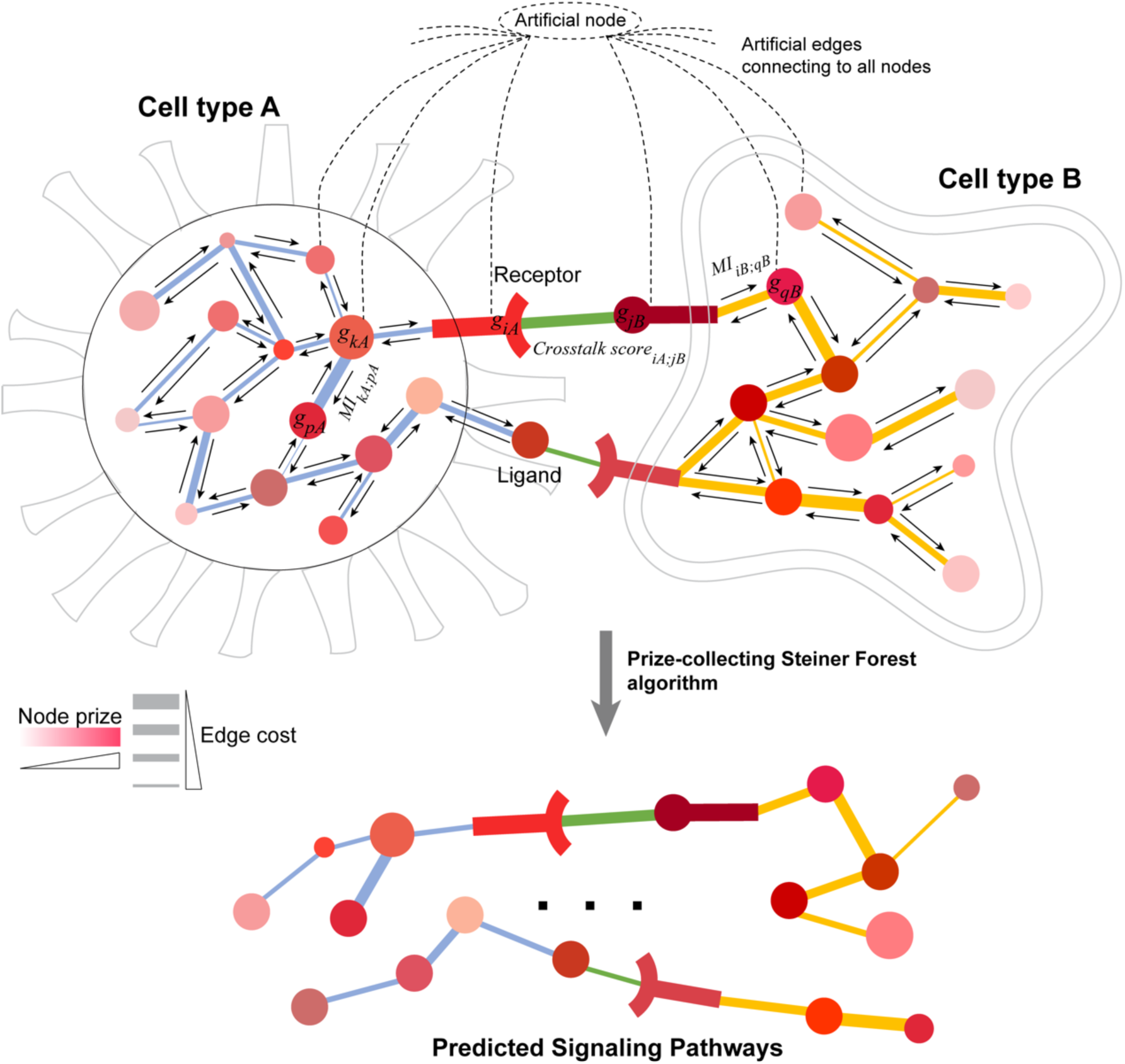
Schematic overview of the CytoTalk algorithm. An integrated gene network is constructed *de novo* using single-cell RNA-Seq data alone. The integrated network consists of two intracellular networks that are connected by known ligand-receptor gene pairs. *g*_*iA*,_ *g*_*jB*_, genes *i* and *j* in cell type *A* and *B*, respectively; *MI*_*kA;pA*_, mutual information between genes *k* and *p* in cell type *A*; *Crosstalk score*_*iA;jB*_, crosstalk score between gene *i* in cell type *A* and gene *j* in cell type *B*. Node color is proportional to node prize and edge thickness is inversely proportional to edge cost. Node prize is computed based on both gene expression specificity and network distance to ligand/receptor gene. Edge cost is computed based on mutual information or crosstalk score. Directed arrows along each edge indicate network propagation procedure. An artificial node (root node) is included in the integrated network to enable search using the prize-collecting Steiner forest (PCSF) algorithm. Statistical significance of the predicted pathways is computed by comparing to null models of PCSFs identified from 1000 randomized networks.

### Performance evaluation using spatial transcriptomics data

To evaluate the performance of CytoTalk, we applied it to a scRNA-Seq data set on mouse visual cortex ^14^. The data set covers the following cell types, glutamatergic neurons, GABAergic neurons, astrocytes, microglia, endothelial cells, oligodendrocytes and oligodendrocyte precursor cells. On average, 6,358 genes were detected per cell. Among the covered cell types, endothelial cells are known to signal to microglia and astrocytes and astrocytes and glutamatergic neurons are known to signal to each other ^15-19^. We identified signaling networks between the three pairs of cell types, endothelial-microglia (EndoMicro), endothelial-astrocyte (EndoAstro) and astrocyte-neuron (AstroNeuro), respectively. The predicted cell-type-specific signaling networks consist of 481, 404, and 1051 genes and involves 51, 44, and 35 ligand-receptor interactions (crosstalk edges), respectively. Compared to PCSFs identified using 1000 randomized input networks, all predicted signaling networks have significantly smaller objective function scores and larger fractions of crosstalk edges (empirical p-values < 0.001, Supplementary Fig. 3). Several predicted ligand-receptor pairs are known to mediate signal transduction between the three cell types. For example, TGFB1 secreted by microglia is known to bind to ACVR11 that is expressed on neighboring endothelial cells in the mouse visual cortex ^20^. Astrocytes are known to express VEGFA that can signal to endothelial cells in the central nervous system (CNS) via KDR (or VEGFR2), which is important for CNS angiogenesis and the formation of the blood-brain barrier ^16,21^. N1GN1 expressed on astrocytes can interact with NRXN1 expressed on neurons to control astrocyte morphogenesis and synaptogenesis ^19^.

To systematically evaluate the performance of CytoTalk, we used a matched sequential fluorescence *in situ* hybridization (SeqFISH+) data set ^20^, covering the same set of cell types as the scRNA-Seq data set. On average, this data set provides spatially resolved abundance of 5,826 transcripts (3,344 genes) per cell. For each pair of cell types under study, we divided cell pairs into three groups based on their physical distances determined using the seqFISH+ data (Supplementary Fig. 1a, 1b). We then calculated the seqFISH+ expression correlation among signaling pathway genes across the three groups of cell pairs (Fig. 3a, Methods). Our rationale is that cells that are close together are more likely to signal to each other. Therefore, *bona fide* signaling pathway genes are expected to have higher spatial expression correlation in these cells than cells that are further apart.

**Fig. 3.**
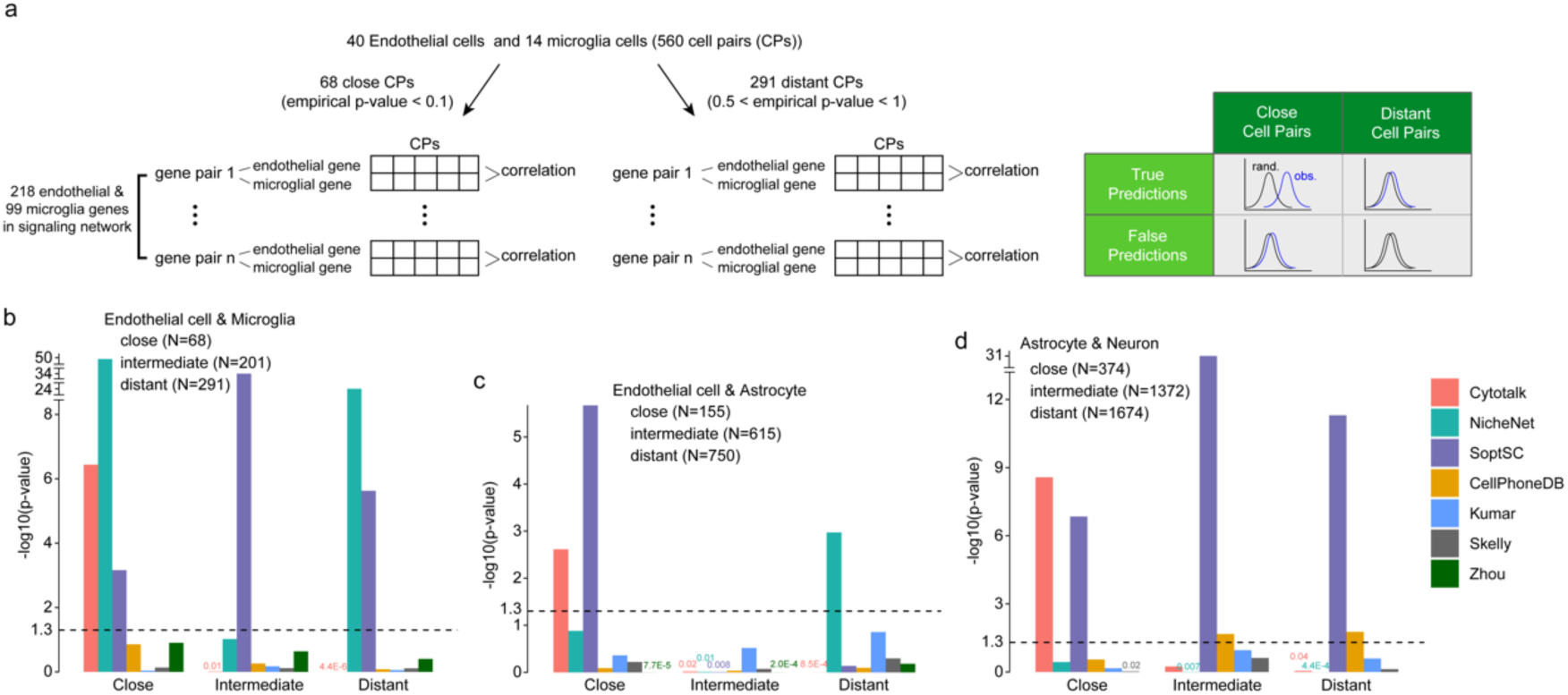
Performance evaluation of the CytoTalk algorithm using spatial transcriptomics data. **(a)** Schematic illustration of the procedure and rationale for using SeqFISH+ data to evaluate predicted signaling networks. **(b-d)** Spatial expression correlation of predicted pathway genes across cell pairs with close, intermediate and far distance. Cell pairs were categorized into three distance groups using SeqFISH+ data. Pearson correlation of pathway genes was computed using SeqFISH+ data and compared to that computed from randomly selected gene pairs. Statistical difference between two distributions of correlation values was computed using one-sided Kolmogorov-Smirnov test for predicted EndoMicro **(b)**, EndoAstro **(c)** and AstroNeuro **(d)** signaling networks.

We compared CytoTalk to six published algorithms, four designed for predicting ligand-receptor pairs ^1-4^ and two designed for predicting full signaling pathways based on known pathway annotations ^7,8^. Since a comprehensive list of true ligand-receptor pairs are not available for the three cell type pairs, we first asked what fractions of the predicted ligand-receptor pairs are shared among the six methods. We reason that a more accurate method will have on average a larger fraction of overlapped predictions with all other methods. Indeed, we found that CytoTalk has the largest average overlap with the rest of the methods (Supplementary Fig. 2), suggesting that CytoTalk has the highest accuracy among the seven methods.

To further benchmark the performance, we next used SeqFISH+ data ^20^ to corroborate pathway predictions using scRNA-Seq data alone ^14^. For endothelial and microglia cells, there are 68, 201, and 291 cell pairs in the close, intermediate and distant groups (Supplementary Fig. 1c). We found that signaling pathways predicted by CytoTalk have significantly larger Pearson correlation coefficients (PCCs) across close endothelial-microglia cell pairs than expected by chance (one-sided Kolmogorov-Smirnov test p-value = 3.6E-7) whereas the PCCs among intermediate and distant cell pairs are indistinguishable from the null distribution, thus providing support to our predictions (Fig. 3a, 3b). In comparison, predicted pathway genes by other methods show no or less significant difference in PCC compared to randomly selected gene pairs among close cell pairs, except for the signaling networks predicted by NicheNet and SoptSC (Fig. 3b). However, pathways predicted by NicheNet and SoptSC also show significantly larger PCCs compared to random gene pairs among intermediate and distant cell pairs, suggesting that those predictions are false positive predictions.

For predicted EndoAstro and AstroNeuro signaling networks, we also found that CytoTalk predictions have significantly larger PCCs *only* among close cell pairs. In comparison, predictions by other methods do not show this trend, except the EndoAstro signaling network predicted by SoptSC (Fig. 3c, 3d). Taken together, these results demonstrate that CytoTalk has significant improvement over published methods.

### Performance evaluation using scRNA-Seq data without receptor gene expression

To further evaluate the accuracy of CytoTalk, we applied it to a scRNA-Seq data set in which the transcriptomes of wild type and receptor gene knockout cells were compared ^22^. The data set covers 13 cell types in the mouse lung, including T and B cells, neutrophils, basophils, monocytes, macrophages, endothelial cells, alveolar type I (AT1) and type II (AT2) cells, club cells, smooth muscle cells, fibroblasts and pericytes. On average, 2,627 transcripts (1,143 genes) were detected per cell. The authors discovered a novel signaling pathway involving interleukin 33 (IL33) secreted by AT2 cells and interleukin 1 receptor like 1 (IL1RL1) on basophils ^22^. Using this data set, we first asked whether the IL33-IL1RL1 interaction between AT2 cells and basophils can be predicted. We found that all three methods (CytoTalk, NicheNet, and SoptSC) can identify the IL33-IL1RL1 interaction. We then evaluated the prediction accuracy using receiver operating characteristic (ROC) curve. To this end, we used the differentially expressed genes (DEGs) between *IL1RL1*-knockout and wild type basophils as the ground truth. We found that predictions by CytoTalk have a higher area under the ROC curve (Fig. 4a). Furthermore, the downstream pathway genes predicted by CytoTalk tend to be more significantly differentially expressed compared to the genes predicted by the other two methods (one-sided Wilcoxon p-values < 1.0E-26, Fig. 4b). Taken together, these results provide additional support for the improved performance of CytoTalk compared to existing methods.

**Fig. 4.**
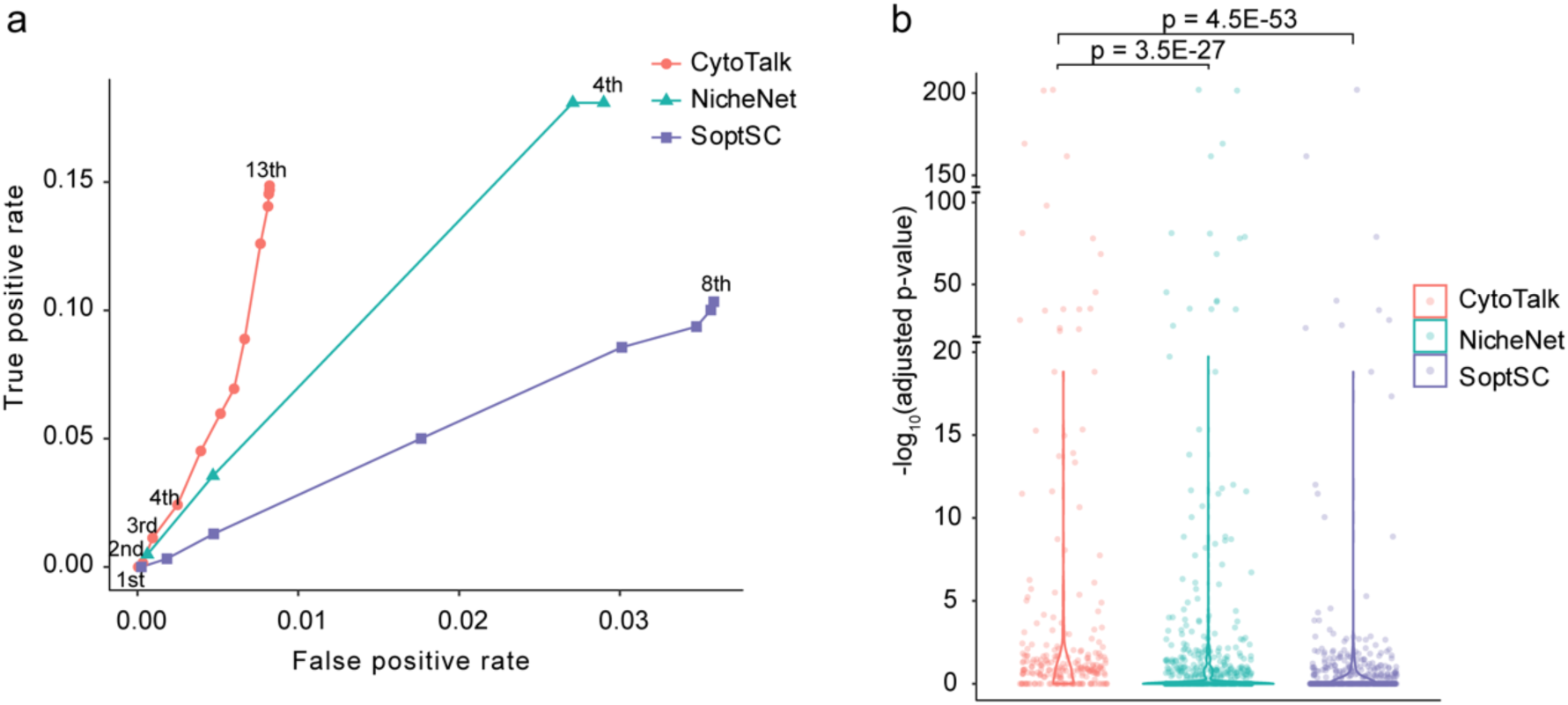
Performance evaluation of the CytoTalk algorithm using scRNA-Seq data without receptor gene expression. Pathways predicted by CytoTalk, NicheNet and SoptSC were evaluated by comparing to differentially expressed genes (DEGs) between receptor gene knockout and wild type cells. DEGs were identified by DESeq2 using a Benjamini-Hochberg (BH) adjusted p-value cutoff of 0.05. **(a)** Receiver operating characteristics curve. DEGs between *IL1RL1*-knockout and wild type basophils are considered true positives. True positive rate and false positive rate were computed using the DEGs. Numbers at each point on the curve indicate the network distance of predicted genes to the receptor (IL1RL1). For instance, 1^st^ means a predicted gene that is a first-order neighbor of the receptor. **(b)** Distribution of p-values of differential expression for all genes in the predicted downstream pathways. Shown are violin plots of -log_10_(BH-adjusted p-value). P-values for comparing distributions were computed using one-sided Wilcoxon rank-sum test.

## Discussion

We introduce a computational method, CytoTalk, for the construction of cell-type-specific signal transduction pathways using scRNA-Seq data. The input to CytoTalk are scRNA-Seq data and known ligand-receptor interactions. Unlike previous methods using known pathway annotations ^7,8^, CytoTalk constructs full pathways *de novo*.

Systematic evaluation of predicted signaling pathways represents another major challenge due to the lack of gold standard pathway annotations. Here, we propose two benchmarking strategies using single-cell spatial transcriptomics data and perturbation-based scRNA-Seq data, respectively. Using spatial transcriptomics data, cells of two types can be stratified into groups of cell-pairs based on the physical distances between them. The predicted signaling pathways can be validated by computing the spatial expression correlation for pathway genes across cell pairs. Using perturbation-based scRNA-Seq data, especially data with ligand/receptor gene knockout, differentially expressed genes between the perturbed and wild type cells can be used to validate predicted pathways. Using these two benchmarking approaches, we demonstrated that CytoTalk significantly outperforms six existing methods that also use scRNA-Seq data to characterize cell-cell communication.

In the current version of CytoTalk, the node prize is defined based on cell-type-specificity of gene expression. Thus, CytoTalk may fail to identify signaling pathways whose genes have low expression specificity in the cell types under study. To address this issue, the node prize can be redefined by considering both absolute gene expression level and cell-type-specificity of gene expression. It is also well known that activity of a signaling pathway is regulated by post-translational modifications. With the rapid development of single-cell proteomics technologies ^23^, CytoTalk can be further improved by incorporating such data.

In summary, CytoTalk provides a much-needed means for *de novo* construction of complete cell-type-specific signaling pathways. Comparative analysis of signaling pathways will lead to a better understanding of cell-cell communication in healthy and diseased tissues.

## Methods

### Construction of intracellular functional gene interaction network

We construct an intracellular gene co-expression network for each cell type by calculating the mutual information between all pairs of genes using the *infotheo* R package ^24^. Edges representing indirect functional relationship between genes are removed using the data processing inequality criterion implemented in the *parmigene* package ^25-27^. Mutual information value is used as the edge weight in the two intracellular networks.

### Crosstalk score of a ligand-receptor pair between two cell types

Cell-cell communication in multi-cellular organism can be mediated by autocrine signaling, paracrine signaling and juxtacrine signaling (contact-mediated signaling). There is a fundamental trade-off between autocrine and paracrine signaling ^28^. The former enables a single cell to talk to itself whereas the latter is designed to allow multiple cell types to talk to each other. Motivated by this observation, we define a crosstalk score between gene *i* in cell type *A* and gene *j* in cell type *B* as below. Gene*s i* and *j* encode a ligand and a receptor or vice versa.

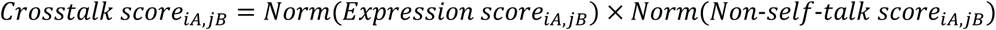

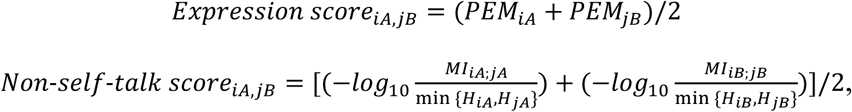

where *Expression score*_*iA,jB*_ is defined as the average preferential expression measure ^29,30^ (*PEM*, defined below) values of gene *i* and *j* in cell types *A* and *B*, respectively. *Non*-*self*-*talk score*_*iA,jB*_ is defined based on information-theoretic measures. *MI*_*iA*;*jA*_ (or *MI*_*iB*;*jB*_) is the mutual information between genes *i* and *j* in cell type *A* (or cell type *B*). min{*H*_*i\**_, *H*_*j\**_} is the upper bound of the mutual information and is used to normalize the mutual information values to [0, 1]. *H*_*iA*_ and *H*_*jA*_ are Shannon entropy of genes *i* and *j* in cell type *A*, respectively. *H*_*iB*_ and *H*_*jB*_ are Shannon entropy of genes *i* and *j* in cell type *B*, respectively. The crosstalk score equals the product of the min-max normalized expression score and non-self-talk score. If genes *i* and *j* are specifically expressed in cell types *A* and *B*, respectively, but are not co-expressed in either cell type (likely involved in self-communication), the *Crosstalk score*_*iA,jB*_ would be high, suggesting a high possibility of crosstalk between the two cell types.

The *PEM* value for defining cell-type-specificity of gene *i* in cell type *A* is defined as following:

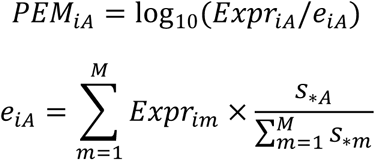

where *Expr*_*iA*_ is the observed expression of gene *i* in cell type *A. e*_*iA*_ is the expected expression of gene *i* in cell type *A* under the null hypothesis of uniform expression across all *M* cell types in the scRNA-Seq data. *Expr*_*i*\*_ represents the expression of gene *i* in cell type *m. s*_*\**\*_ is the sum of expression of all genes in cell type *m. s*_*\*A*_ is the sum of expression of all genes in cell type *A*. Since we focus on genes that are expressed higher in a cell type rather than lower, *PEM*_*iA*_ is set to zero if it is negative.

### Construction of an integrated network between two cell types

We construct an integrated network consisting of two intracellular networks connected by known ligand-receptor interactions. We collected 1,941 manually annotated ligand-receptor interactions, including 1,894 interactions from ^31^ and 47 interactions from ^32-36^ (Supplementary Table 2). Note that both secreted and cell-surface proteins could be ligands. For each ligand-receptor pair, if the ligand gene and the receptor gene are present in the two intracellular networks, we connect them and denote the edge as a crosstalk edge. The crosstalk score is used as the edge weight as described above. Due to the difference in scale between mutual information value and crosstalk score, we separately normalize the edge weights of intracellular networks and crosstalk edges using z-score transformation.

### *De novo* identification of signaling network between two cell types

We formulate the identification of a signaling network between two cell types as a prize-collecting Steiner forest (PCSF) problem ^12,37^. Because the forest is a disjoint set of trees, PCSF problem is a generalization of the classical prize-collecting Steiner tree (PCST) problem ^38,39^. The individual signaling pathways are represented as trees, the collection of which (forest) represents the entire signaling network between two cell types.

We define edge costs and node prizes in the integrated network as follows. The z-score normalized edge weights of the integrated network are first scaled to the range of [0, 1]. Edge cost is then defined as 1 − *scaled*_*edge*_*weight*. Node prize is defined based on both *PEM* value of a gene and its closeness to the ligand/receptor genes in the network in order to identify signaling networks centered around the crosstalk edges. To capture the closeness, we use a network propagation procedure to calculate a relevance coefficient for each gene in an intracellular network.

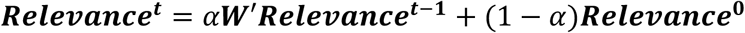

where ***Relevance***^***t***^ is the relevance coefficient vector for all genes in the intracellular network at iteration *t*. ***Relevance***^**0**^ is the initial value of the relevance coefficient vector such that *Relevance*^2^(*i*) = 1 if gene *i* is a ligand or receptor. Otherwise, *Relevance*^2^(*i*) = 0. ***W***′ is a normalized edge weight matrix for an intracellular network, which is defined as ***W***′ = ***D***^−**1**/**2**^***WD***^−**1**/**2**^. Here, ***W*** is set to the original mutual information matrix and ***D*** is defined as a diagonal matrix such that *D*(*i, i*) is the sum of row *i* of the matrix ***W***. This network propagation procedure is equivalent to a random walk with restart on the network. *α* is a tuning parameter that controls the balance between prior information (known ligands or receptors) and network smoothing. Node prize of a gene is defined as the product of its *PEM* value and the relevance coefficient to capture both the cell-type-specificity and the closeness of this gene to the ligand or receptor gene in the network. To avoid extremely large node prizes for ligand or receptor genes, we used *α* = 0.9 in this study.

The PCSF algorithm identifies an optimal forest in a network that maximizes the total amount of node prizes and minimizes the total amount of edge costs in the forest. While PCSF problem is NP-hard and often needs a high computational cost ^37^, we employ a PCSF formulation established in ^37,40^ and use a highly efficient prize-collecting Steiner tree (PCST) algorithm ^41,42^ to identify the PCSF. The objective function of the PCSF problem is defined as below.

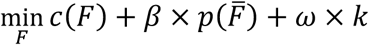

where *F* represents a forest (i.e. multiple disconnected trees) in the integrated network. *c*(*F*) denotes the sum of edge costs in the forest *F* and 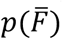 denotes the sum of node prizes of the remaining subnetwork excluding the forest *F* from the network. We modify the integrated network by introducing an artificial node and a number of artificial edges to the original network. The artificial edges connect the artificial node to all genes in the original network. The costs of all artificial edges are the same and are defined as *ω*, which influences the number of trees, *k*, in the resulting PCSF. *β* is a parameter for balancing the edge costs and node prizes, which influences the size of the resulting PCSF. By tuning parameters *β* and *ω*, multiple PCSTs can be identified with the artificial node as the root node. For each identified PCST, a PCSF can be obtained by removing the artificial node and artificial edges from the PCST.

We identify the signaling network between two cell types by searching for a robust PCSF across the full parameter space (Supplementary Fig. 4). For each identified PCSF, we compute the occurrence of each edge in all identified PCSFs to construct a background distribution of edge occurrence frequency. Next, we calculate a p-value for each PCSF by comparing the edge occurrence frequency distribution of this PCSF to the distribution of all other identified PCSFs using one-sided Kolmogorov-Smirnov test. The PCSF with the minimum p-value is considered as the most robust signaling network predicted by CytoTalk.

To further evaluate the statistical significance of the identified PCSF, we construct null distributions for the objective function and for the fraction of crosstalk edges in a PCSF using 1000 null PCSFs identified from randomized integrated networks (Supplementary Fig. 3). To generated the randomized networks, we separately shuffle the edges of the two intracellular networks while preserving the node degree distribution, node prizes and crosstalk edges as the original integrated network.

### Parameter selection

The main parameters of CytoTalk are *β* and *ω* in the objective function of the PCSF problem. We first determined the optimal ranges of the two parameters based on the size and overlap of the resulting PCSFs. For the mouse visual cortex data set, *β* values were tested from 1 to 60 with a step size of 1. For the mouse lung tissue data set, *β* values were tested from 5 to 300 with a step size of 5. We found using *β* values above these ranges resulting in very large PCSFs (> 2000 edges) (Supplementary Fig. 4a-4d). For both data sets, *ω* values were tested from 0.1 to 1.5 with a step size of 0.1. We found using *ω* values above this range resulting in PCSFs with little difference compared to existing PCSFs (Supplementary Fig. 4e-4h). Subsequent optimal parameter selection was conducted using the above parameter ranges. For all PCSFs identified using the *β* and *ω* ranges determined above, the occurrence frequency of each edge in a PCSF was computed to construct a background distribution of edge occurrence frequency. A p-value for each PCSF was computed by comparing the edge occurrence frequency distribution of this PCSF to the distribution of all other PCSFs using one-sided Kolmogorov-Smirnov test. The PCSF with the minimum p-value (red dot) was considered as the most robust signaling network predicted by CytoTalk (Supplementary Fig. 4i-4l).

### Processing of scRNA-Seq data

For all scRNA-Seq data sets used in this study (Supplementary Table 1), we only retained protein-coding genes based on annotations from the GENCODE database ^43^. We removed genes expressed in less than 10% of all cells of a given type.

For identifying differentially expressed genes between *IL1RL1*-knockout and wild type basophils, we first filtered out genes that have fewer than 5 sequencing counts in at least 5 cells. Then, we used the *zinbwave* function in the *zinbwave* R package ^44^ to model the zero inflation of the counts. The *DESeq2* R package ^45^ was used to perform differential expression analysis. P-values were adjusted for multiple testing using the method of Benjamini-Hochberg ^46^. 619 differentially expressed genes with adjusted p-values < 0.05 were identified.

### Processing of SeqFISH+ data

The SeqFISH+ data set was downloaded from a published study ^20^, including a cell by gene count matrix with 523 visual cortex cells as rows and 10,000 genes as columns, cell type annotation and cell spatial location (two-dimensional coordinates) data (Supplementary Table 3). Based on the authors’ preprocessing procedure ^20^, we first log2-transformed the count matrix followed by z-score transformation.

### Quantification of expression correlation of signaling pathway genes using SeqFISH+ data

Cell pairs consisting of two types were categorized into three groups based on their physical distance measured by SeqFISH+, namely close, intermediate and distant groups (Supplementary Fig. 1a). The distance cutoffs for the three groups were determined using empirical p-values of 0.1, 0.5 and 1.0, respectively, based on a null distribution of distances between 10,000 randomly selected cell pairs from all cells profiled by SeqFISH+ (Supplementary Fig. 1b-e). Genes of predicted signaling pathways were intersected with the gene set detected by SeqFISH+. Among these genes, we computed a Pearson correlation coefficient of SeqFISH+ expression values between any gene pair (one from each cell type and are connected in the predicted signaling pathways) across individual cells of the two types.

### Running of published methods

NicheNet ^8^ uses a prior ligand-target regulation model by integrating known signaling transduction and transcriptional regulatory interactions to predict a signaling network between two cell types. Given scRNA-Seq data of cell type *A* and *B*, we first defined two gene sets that are expressed in these two cell types, respectively. We used the same expression cutoff and only retained protein-coding genes for analysis as described above. NicheNet requires a pre-defined gene set of interest in cell type *B* as candidate target genes regulated by ligands in cell type *A*. This gene set was defined as the genes that are specifically expressed (i.e. *PEM* score > 0) in cell type *B*. Using these gene sets as the input, we predicted ligands, their signaling pathways, and target genes using *predict_ligand_activities* and *get_ligand_signaling_path* functions in the *NicheNet* R package. These identified signaling pathways are considered as a “cell type *A* to *B*” signaling subnetwork. Using another set of genes that are specifically expressed in cell type *A* as input, we also predicted a “cell type *B* to *A*” signaling subnetwork. By combining the two subnetworks, we obtained a final predicted signaling network between the two cell types.

SoptSC ^7^ also uses known pathway annotations to predict signaling pathways between two cell types. For comparison, we used mouse Reactome pathways and the same ligand-receptor pairs as used by CytoTalk. For each ligand-receptor pair, we computed a matrix of signaling probabilities between any two cells from cell types *A* and *B* using the *LR_Interaction* function in the *SoptSC* MATLAB package. Using this matrix, we computed two probabilities of signaling via the given ligand-receptor pair from each direction (*A* → *B* and *B* → *A*), respectively. Based on these signaling probabilities, we selected the top 10% of ligand-receptor pairs and their known downstream pathways for performance comparison.

Different from NicheNet and SoptSC, the other four methods only predict active ligand-receptor pairs between two cell types. Among these four methods, Zhou’s and Skelly’s methods are similar, which consider the gene expression levels of a ligand and its receptor separately. Based on Zhou’s method ^3^, for each gene *i*, we calculated the mean 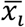 and standard deviation σ_*i*_ of the gene expression values across all cell types. If the average expression values of the ligand gene in cell type A and the receptor gene in cell type B are both larger than 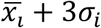, this ligand-receptor pair is predicted to be active by Zhou’s method. Based on Skelly’s method ^1^, for each ligand-receptor pair, if the ligand and the receptor genes are expressed in more than 20% of the cells of cell types *A* and *B*, respectively, this ligand-receptor pair is retained and considered to transmit a signal from cell type *A* to *B*. We considered all retained ligand-receptor pairs from both directions (*A* → *B* and *B* → *A*) as the final predictions by Skelly’s method.

Kumar’s method ^2^ is different from the two methods above, which defines an interaction score for a given ligand-receptor pair as the product of the average expression of the ligand gene in cell type *A* and the average expression of the receptor gene in cell type *B*. We selected the top 10% of ligand-receptor pairs based on interaction scores as the final predictions by Kumar’s method.

CellPhoneDB ^4^ is a repository of curated ligand-receptor pairs and can be used for predicting ligand-receptor interactions based on their cell-type specificity. Given a scRNA-Seq gene expression matrix and cell type annotation data as the input, we used the *cellphonedb* function in the *CellPhoneDB* Python package to compute a p-value for the likelihood of cell-type specificity of each ligand-receptor pair. For two given cell types *A* and *B*, we selected the ligand-receptor pairs with p-values < 0.05 from each direction (*A* → *B* and *B* → *A*) as the final predictions by CellPhoneDB.

## Supplementary Figures

**Supplementary Fig. 1.**
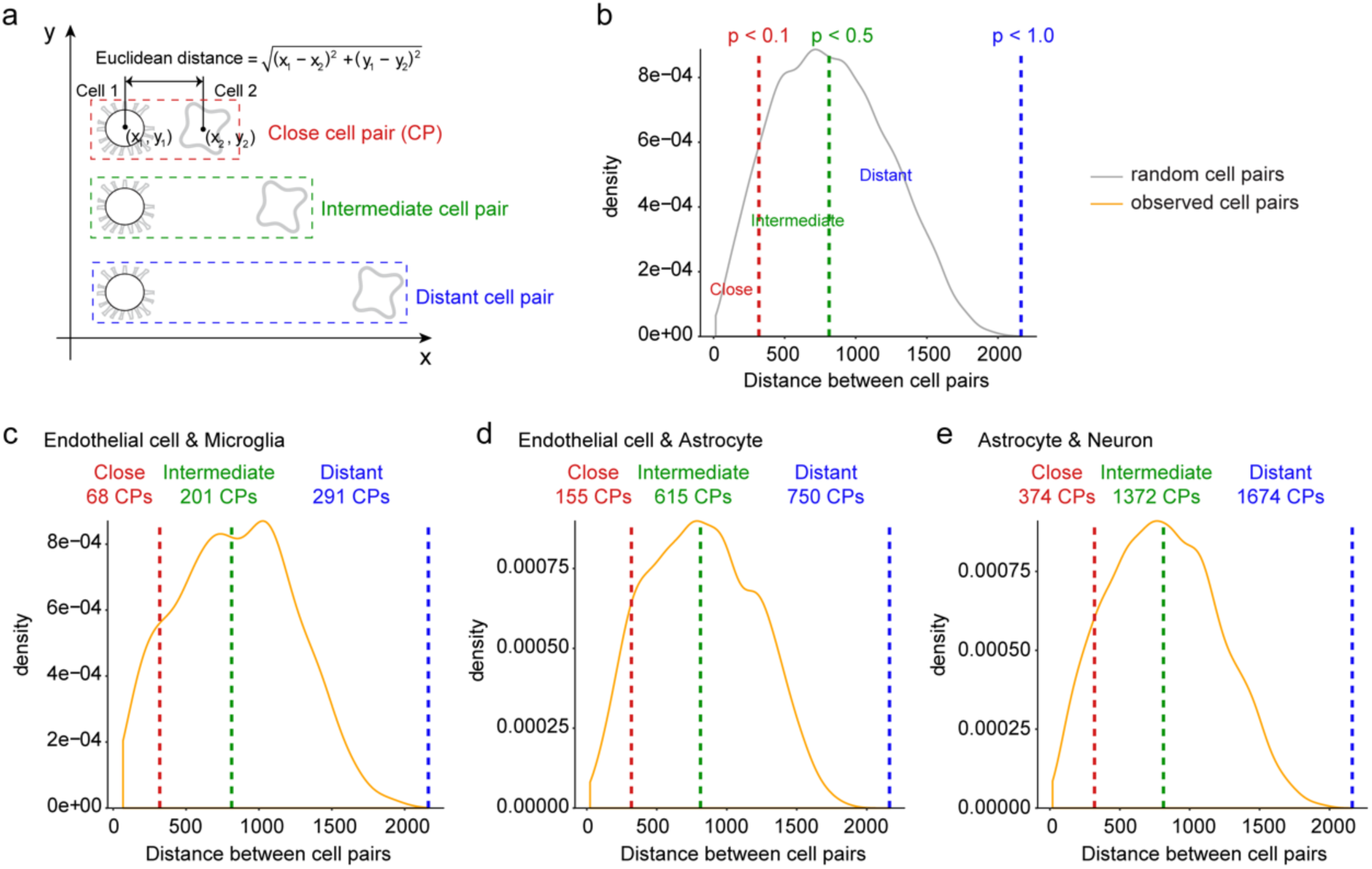
Stratification of cell pairs based on their physical distance in the SeqFISH+ data. **(a)** Each cell profiled by SeqFISH+ has a spatial coordinate (x, y). Cell pairs of two different types were grouped into three groups based on their Euclidean distances computed using their coordinates, namely close, intermediate and far distance groups. **(b)** The distance cutoffs for the three groups were determined using empirical p-values of 0.1, 0.5 and 1.0, respectively, based on a null distribution of distances between 10,000 randomly selected cell pairs in the SeqFISH+ data set. Using these distance cutoffs, 560, 1520 and 3420 cell pairs were divided into three groups for endothelial and microglial cells (EndoMicro) **(c)**, endothelial and astrocytes (EndoAstro) **(d)** and astrocytes and neurons (AstroNeuro) **(e)**, respectively.

**Supplementary Fig. 2.**
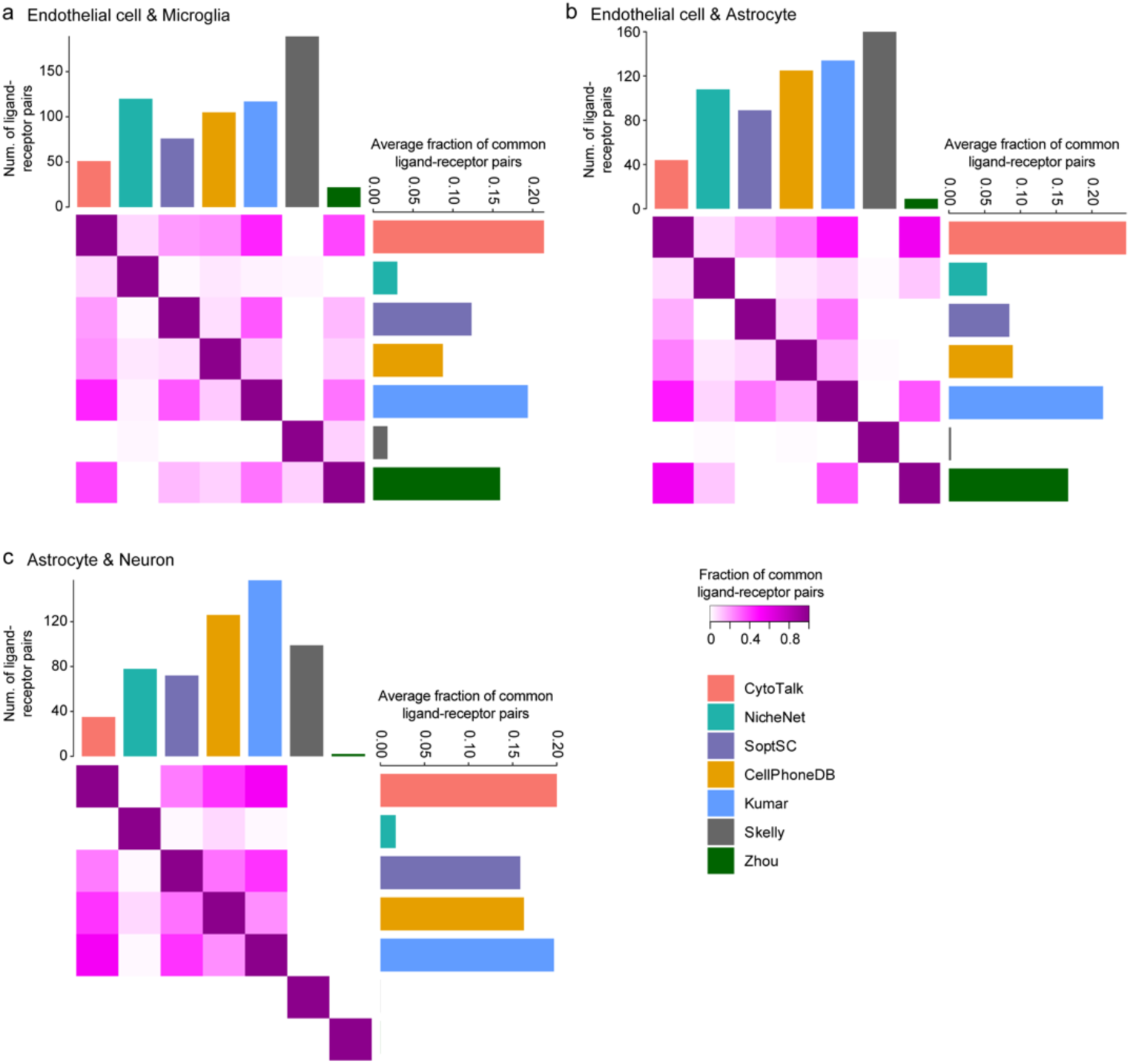
Comparison of ligand-receptor pairs predicted by seven methods. Ligand-receptor pairs were predicted by seven methods for EndoMicro **(a)**, EndoAstro **(b)** and AstroNeuro **(c)**. The overlap fractions of ligand-receptor pairs between any two methods are shown in the heatmap. Shade of magenta is proportional to the overlap fraction, which is defined as the number of overlapped ligand-receptor pairs divided by the minimum number of identified ligand-receptor pairs between two methods. The number of ligand-receptor pairs predicted by each method is shown in the bar plot above the heatmap. The average overlap fraction between a given method and the other methods is shown in the bar plot to the right of the heatmap.

**Supplementary Fig. 3.**
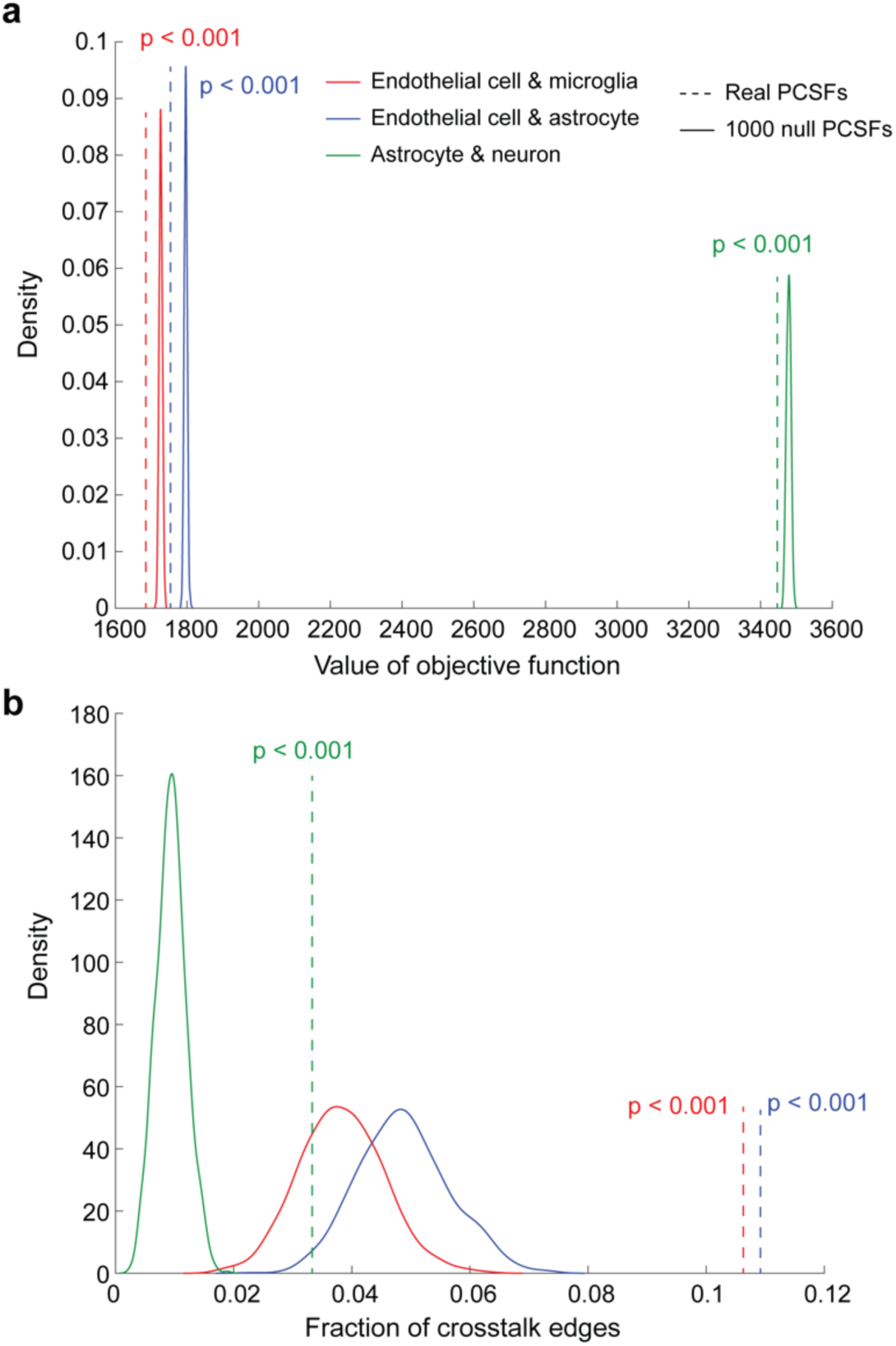
Statistical significance of predicted signaling networks. Statistical significance of predicted signaling networks were evaluated based on the objective function of the prize-collecting Steiner forest (PCSF) algorithm **(a)** and the fraction of crosstalk edges in the PCSF **(b)**. Empirical p-value was calculated by comparing the PCSF identified from the real network (dashed line) to PCSFs identified from 1000 randomized networks (solid line). To generate the randomized networks, given the real network, we separately shuffled the edges of the two intracellular networks keeping the node degree distribution, node prizes and crosstalk edges the same as the real network. Then, these randomized integrated networks together with the same *β* and *ω* values as the predicted signaling network were used as inputs for the PCSF algorithm to generate null models of PCSFs.

**Supplementary Fig. 4.**
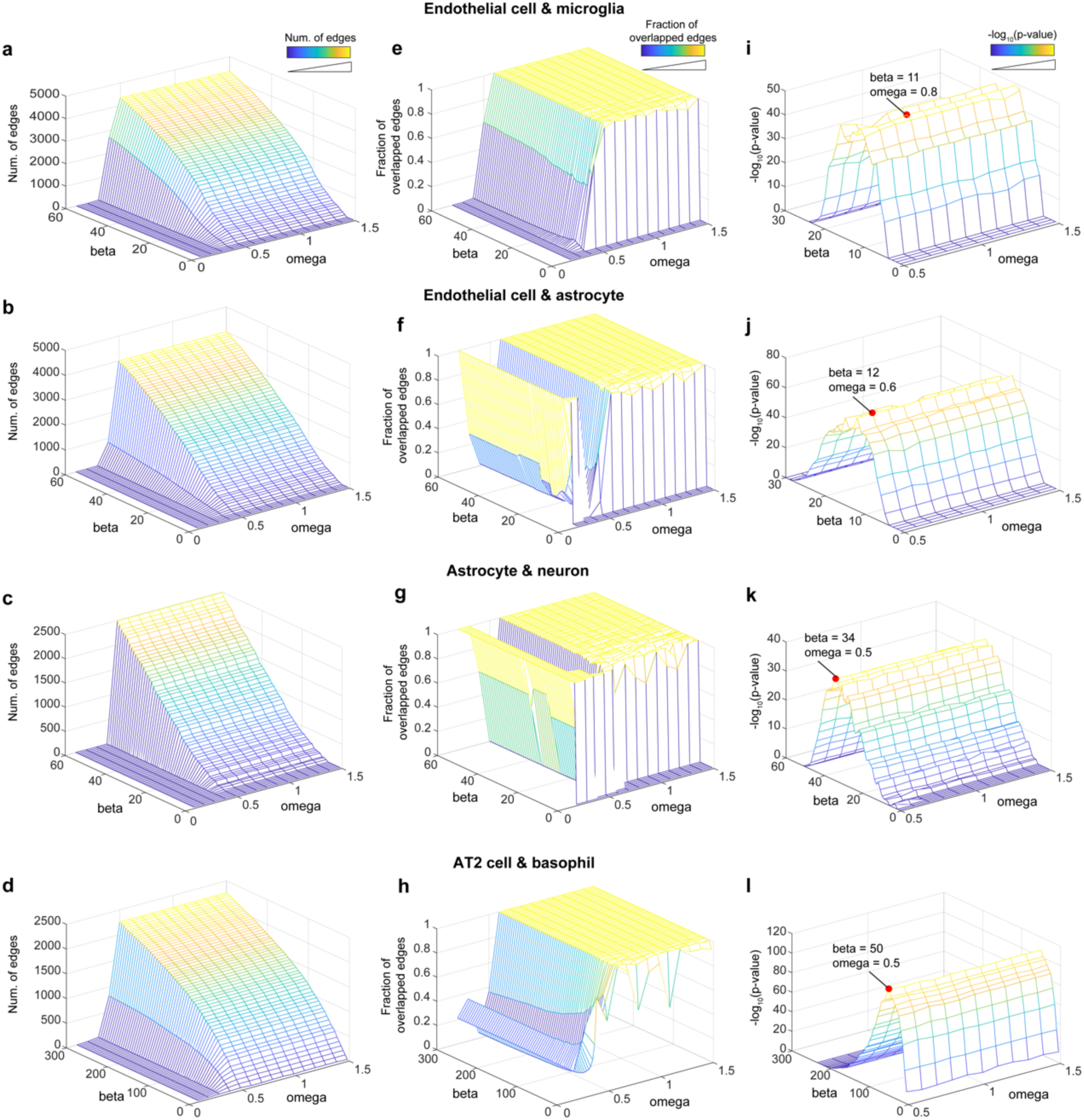
Parameter selection for the CytoTalk algorithm. The optimal ranges of two main parameters of the algorithm, *β, ω*, were determined first. Ranges of *β* and *ω* values were tested for the mouse visual cortex and lung data sets. The number of edges in these PCSFs **(a-d)** and the fraction of overlapped edges between PCSFs using adjacent omega values **(e-h)** are shown. **(i-l)** Determination of parameter settings that result in most robust PCSF. For all PCSFs identified using the *β* and *ω* ranges determined above, the occurrence frequency of each edge in a PCSF was computed to construct a background distribution of edge occurrence frequency. A p-value for each PCSF was computed by comparing the edge occurrence frequency distribution of this PCSF to the distribution of all other PCSFs using one-sided Kolmogorov-Smirnov test. The PCSF with the minimum p-value (red dot) was considered as the most robust signaling network predicted by CytoTalk. All 3-D mesh surface plots use *z*-axis values for both height and color.

## Supplementary Tables

**Supplementary Table 1.** Summary of data sets used in this study.

| Data set | Accession number/URL | PubMed ID |
| --- | --- | --- |
| Mouse visual cortex<br>scRNA-Seq | GSE71585 | 26727548 |
| Mouse visual cortex<br>(SeqFISH+) | https://github.com/CaiGroup/seqFISH-PLUS | 30911168 |
| Mouse mammary<br>gland | GSE109774 | 30283141 |
| Mouse skin | GSE109774 | 30283141 |
| Mouse lung | GSE19228 | 30318149 |

**Supplementary Table 2. List of known ligand-receptor interactions used in this study.**

**Supplementary Table 3. SeqFISH+ data used in the performance evaluation.** The table includes cell type annotation and spatial location (two-dimensional coordinates) data of 523 visual cortex cells. A 10,000 gene count matrix of these cells are also appended.

